# The genetic Allee effect: A unified framework for the genetics and demography of small populations

**DOI:** 10.1101/038125

**Authors:** Gloria M. Lucque, Chloé Vayssade, Benoît Facon, Thomas Guillemaud, Franck Courchamp, Xavier Fauvergue

## Abstract

The Allee effect is a theoretical model predicting low growth rates and the possible extinction of small populations. Historically, studies of the Allee effect have focused on demography. As a result, underlying processes other than the direct effect of population density on fitness components are not generally taken into account. There has been heated debate about the potential of genetic processes to drive small populations to extinction, but recent studies have shown that such processes clearly impact small populations over short time scales, and some may generate Allee effects. However, as opposed to the ecological Allee effect, which is underpinned by cooperative interactions between individuals, genetically driven Allee effects require a change in genetic structure to link the decline in population size with a decrease in fitness components. We therefore define the genetic Allee effect as a two-step process whereby a decrease in population size leads to a change in population genetic structure, and in turn, to a decrease in individual fitness. We describe potential underlying mechanisms, and review the evidence for this original type of component Allee effect, using published examples from both plants and animals. The possibility of considering demogenetic feedback in light of genetic Allee effects clarifies the analysis and interpretation of demographic and genetic processes, and the interplay between them, in small populations.

## Introduction

The Allee effect can be defined as a decrease in fitness caused by a decrease in population size (Stephens et al. 1999). The Allee effect jeopardizes the persistence of small populations, whether declining or bottlenecked (threatened species; Angulo et al. 2007, Bonsall et al. 2014, Kuparinen et al. 2014); (introduced/invasive species; Davis et al. 2004, Taylor and Hastings 2005, Johnson et al. 2006). This concept, first introduced by the American ecologist Warder Clyde Allee in the 1930s, has been the focus of growing interest from population ecologists over the last two decades. In 1999, Stephens *et al.* formalized the definition of Allee effects, making a crucial distinction between component and demographic Allee effects. A component Allee effect is a decrease in the value of any component of individual fitness caused by a decrease in population size. A demographic Allee effect is the consequent decrease in the *per capita* growth rate of the population caused by a decrease in population size and resulting from one or several component Allee effects. Allee effects have been discovered in a wide range of taxa, with various mechanisms underpinned by a shortage of cooperative interactions at low density (Courchamp et al. 1999, Courchamp et al. 2008).

Population dynamicists working on the Allee effect may have overlooked the fact that genetic processes can also impact populations at an ecological time scale of only a few generations (Glémin 2003, Spielman et al. 2004). This is particularly relevant when genetic processes are combined with demographic processes, such as demographic and environmental stochasticity (Tanaka 2000, Hanski and Saccheri 2006). Some of these genetic mechanisms can generate component Allee effects (Courchamp et al. 1999) because they generate a positive relationship between population size and fitness (Fischer et al. 2000, Willi et al. 2005, Berec et al. 2007). These mechanisms are inbreeding depression (Frankham 1995, Spielman et al. 2004), loss of genetic variation (Gomulkiewicz and Holt 1995, Amos and Balmford 2001), and the accumulation of deleterious mutations (Lande 1994, Lynch et al. 1995b).

Contrasting with the pervasive evidence that genetic mechanisms affect the dynamics of small populations, even in the short term, fewer than ten publications have made specific reference to a genetic Allee effect. There are two possible reasons for the lack of studies considering genetic Allee effects. First, although mentioned in a handful of studies, the genetic Allee effect has never been formally defined, so the concept may still be too vague to stimulate new studies. Second, the genetic mechanisms occurring in small populations are the classic territory of population genetics, whereas the Allee effect is a concept derived principally from population dynamics. These two communities view populations differently and usually work on different entities (*e.g.*, individual numbers *vs.* genotype frequencies), concepts and time scales. Most studies in population genetics assume populations of constant size, unaffected by individual fitness (even when modeling inbreeding depression; e.g., Glémin 2003) and dynamicists usually view genetic processes as affecting populations only in the long term. This would make them likely to think that small populations do not generally stay small for long enough (because they grow or disappear due to demographic processes) to suffer from genetic Allee effects. There may have been too little dialog between the two disciplines as yet (Kokko and Lopez-Sepulcre 2007, Metcalf and Pavard 2007, Pelletier et al. 2009) for the genetic Allee effect to have emerged as a robust unifying paradigm.

The aim of this work is to propose the genetic Allee effect as a heuristic framework for studying the interplay between genetics and demography in small populations. We first propose a formal definition for the genetic Allee effect and describe the processes potentially underlying this effect. We review the evidence for genetic Allee effects, including population genetic studies, which, although not referring to the Allee effect, provide numerous examples of genetic Allee effects. Based on these examples, we then propose methods for detecting genetic Allee effects and distinguishing them from ecological Allee effects. In the last section of the paper, we discuss the specificity, limits, and demographic consequences of genetic Allee effects.

## Definition

One fundamental principle of evolutionary biology is that a population is a collection of different genotypes vehicled by a number of different individuals. Changes in genotype frequencies correspond to changes in the number, frequency and association of different alleles within and between loci. These genotype frequencies are referred to hereafter as the (within-population) genetic structure. Changes in genetic structure yield changes in the fitness components that these genotypes influence (Soulé 1980). This relationship underlies the genetic Allee effect because population size is one of the determinants of genetic structure (Nielsen and Slatkin 2013). Here, (following the evolutionary paradigm described by Waples and Gaggiotti 2006), the term “population” refers to a “a group of individuals of the same species living in close enough proximity that any member of the group can potentially mate with any other member”. Unlike population dynamicists, we use the term “population size” to mean the number of individuals that effectively reproduce. We prefer to use this term, rather than effective population size, because the latter is difficult to measure in natural populations and has an ambiguous meaning resulting from its various definitions (the “inbreeding effective population size”, the “variance effective population size”, and the “eigenvalue effective population size”; see details in Sjödin et al. 2005). Population size, genetic structure, and fitness components are the three cornerstones of the genetic Allee effect.

The term “genetic Allee effect” is not new (first appearance in Fischer et al. 2000), but it has seldom been used, and still lacks a proper definition. We define the genetic Allee effect as a two-step process characterized by (1) a change in population genetic structure due to a decrease in population size, and (2) a consequent decrease in individual fitness. This decrease in individual fitness may subsequently lead to a further decrease in population size. These two steps act sequentially to generate a genetically driven component Allee effect. The first step of the genetic Allee effect occurs when population size affects at least one of the following aspects of genetic structure: heterozygosity, frequency of beneficial or detrimental alleles, or allelic richness (Fig. 1). During the second step, any of these changes in population genetic structure may decrease a component of individual fitness through inbreeding depression, drift load or migration load. If, and only if, both steps occur, a genetic Allee effect occurs. The genetic Allee effect therefore contrasts with what could be referred to as an ecological Allee effect, in which individual fitness decreases as a straightforward, one-step consequence of a decrease in population size or population density.

**Figure 1.**
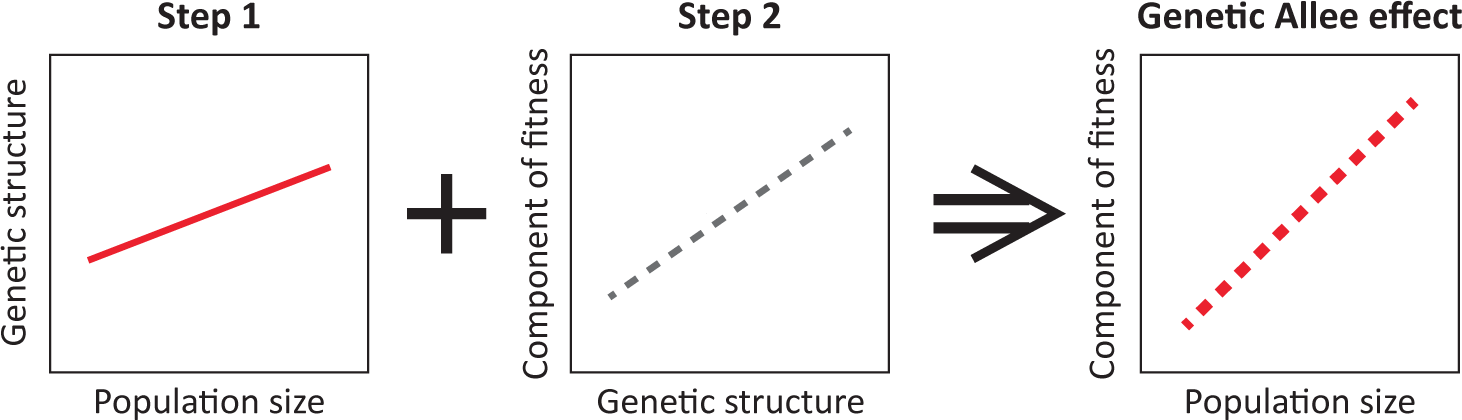
Two successive steps are necessary for a genetic Allee effect to occur. Step 1: a decrease in population size *causes* a change in (a parameter of) genetic structure (heterozygosity, frequency of detrimental and beneficial alleles, allelic richness). Step 2: The change in genetic structure *causes* a decrease in the value of a component of fitness through inbreeding depression, drift load or migration load. When both steps occur in a population, a component genetic Allee effect occurs (left panel).

## Mechanisms underlying genetic Allee effects

Our literature search identified 15 studies showing strong evidence for one or several genetic Allee effects in natural or experimental populations (Table 1). Some of these studies did not actually use the term genetic Allee effect. We also identified about another 40 studies suggesting the occurrence of genetic Allee effects (see for instance the literature cited in Leimu et al. 2006). Following a thorough analysis of these published studies, we defined three types of genetic mechanisms generating Allee effects, each involving a major evolutionary force: inbreeding, drift, or migration.

**Table 1.**
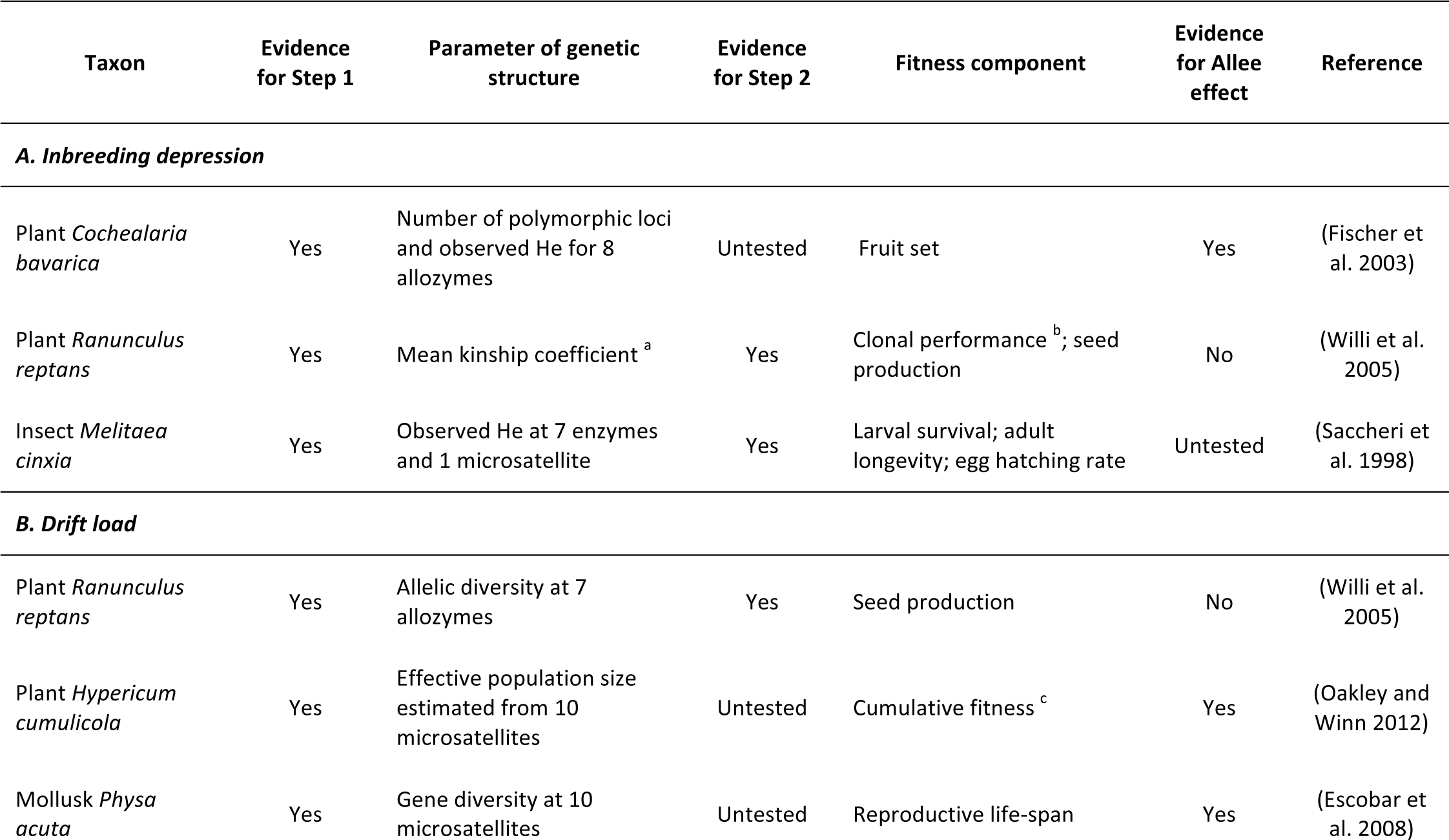
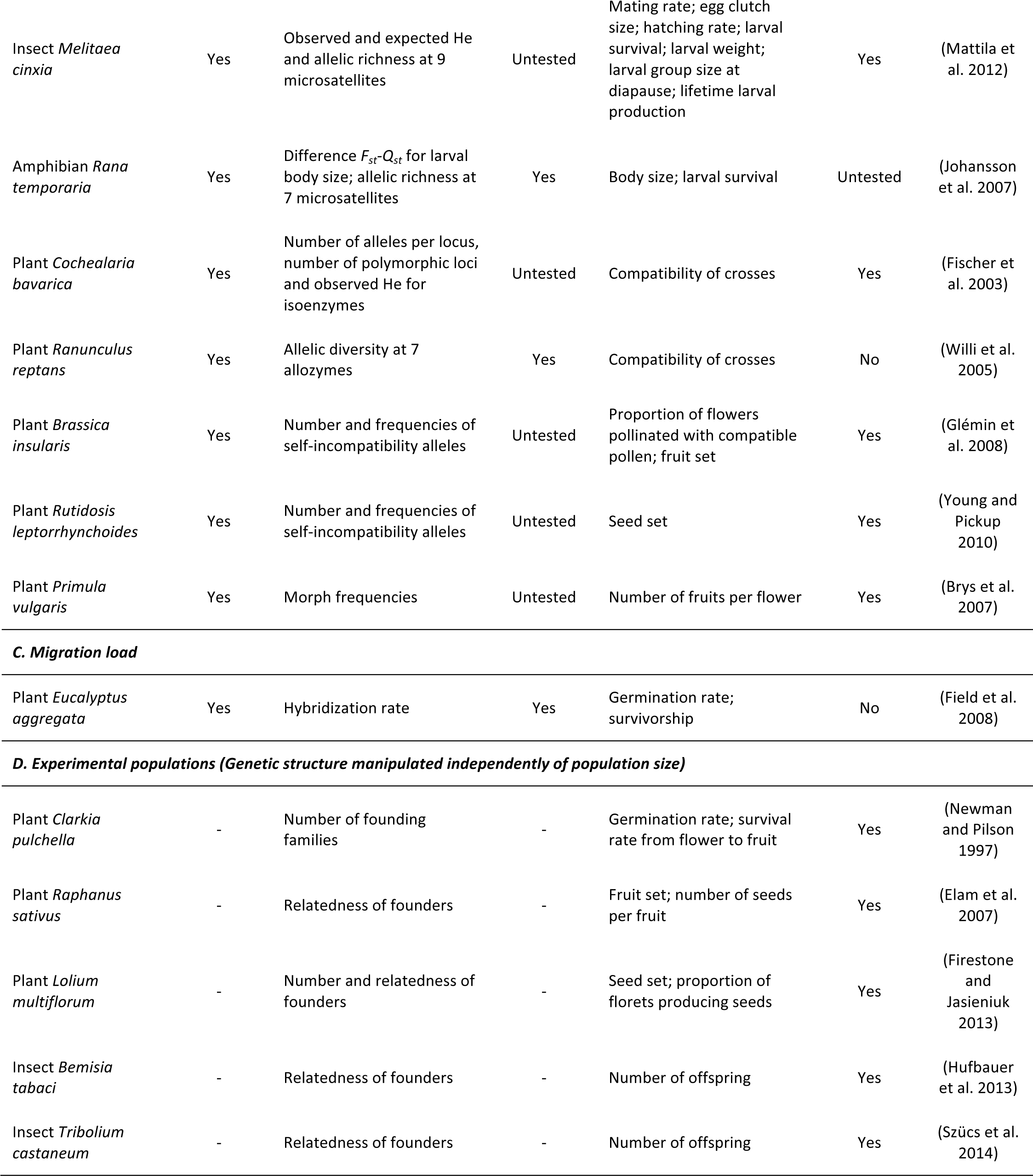
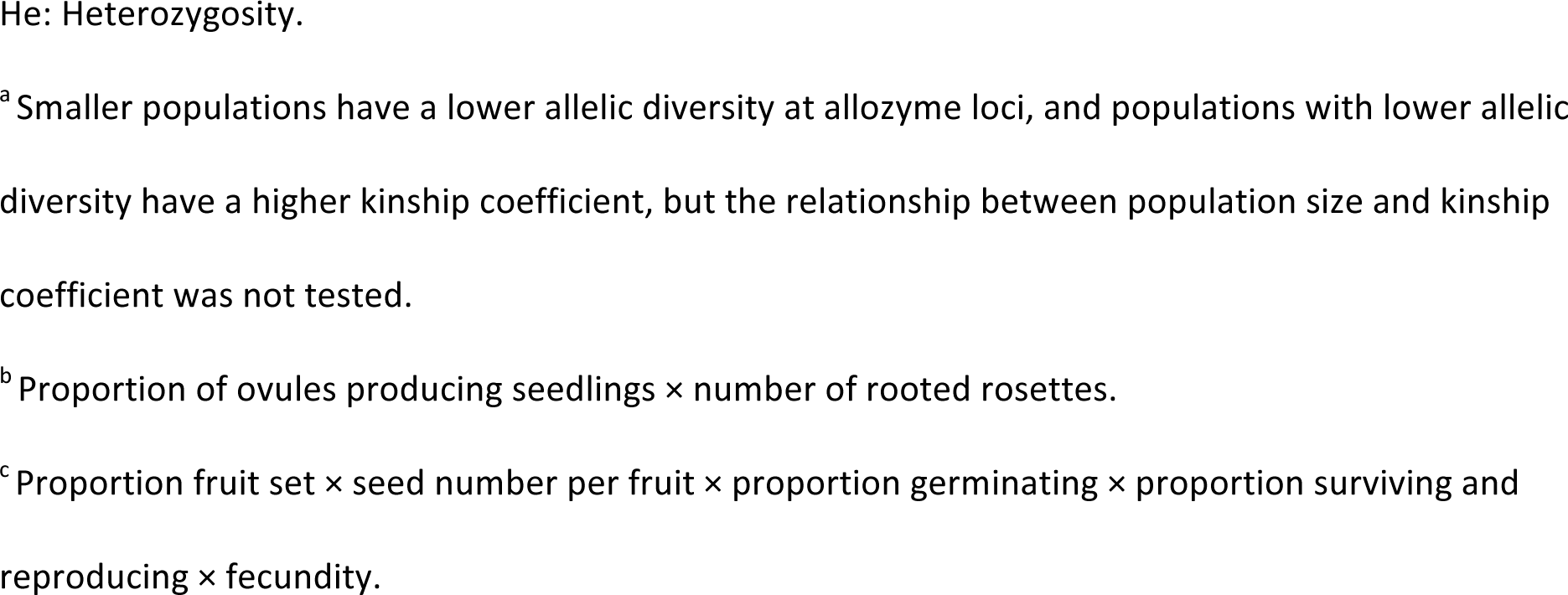
Published evidence for genetic Allee effects in natural and experimental populations. Genetic Allee effects in natural populations are classified according to their underlying mechanism. Evidence for Step 1 indicates whether a relationship between population size and genetic structure was tested and found in the expected direction. Evidence for Step 2 indicates whether a relationship between genetic structure and a component of fitness was tested and found in the expected direction. Strong evidence for a genetic Allee effect implies evidence for both Steps 1 and 2. Evidence for Allee effect indicates whether a positive relationship between population size and a fitness component was observed; this corroborates but does not prove that a genetic Allee effect is occurring. Experimental populations are populations in which genetic structure was manipulated independently of population size. For the four examples concerning experimental populations, the mechanism causing the genetic Allee effect was not identified.

### Inbreeding depression

As population size declines, inbreeding becomes more frequent (Fig. 2; Malécot 1969). An inbred individual results from a cross between two genetically related individuals, and is characterized by a high inbreeding coefficient. Inbred individuals have fewer heterozygous loci than outbred individuals, resulting in a decrease in heterozygosity in declining populations (Frankham 1996, 1998). A pervasive consequence of low heterozygosity is inbreeding depression, defined as the lower fitness of inbred than of outbred individuals. Hence, inbreeding followed by inbreeding depression fulfills the two conditions defining the genetic Allee effect: (1) a decrease in population size causing a change in population genetic structure (a decrease in heterozygosity) and (2) a decrease in one or several components of fitness caused by this change (Fig. 2; Frankham 1998, Reed and Frankham 2003, Spielman et al. 2004). The mechanisms causing inbreeding depression (increased expression of deleterious recessive alleles, overdominance, and epistasis; Li et al. 2008, Charlesworth and Willis 2009) underlie the second step of a genetic Allee effect. For instance, the buttercup *Ranunculus reptans* suffers from a genetic Allee effect due to inbreeding depression (Table 1A; Willi et al. 2005): small populations have a higher mean inbreeding coefficient (Step 1) and populations with higher inbreeding coefficients have lower levels of seed production (Step 2).

**Figure 2.**
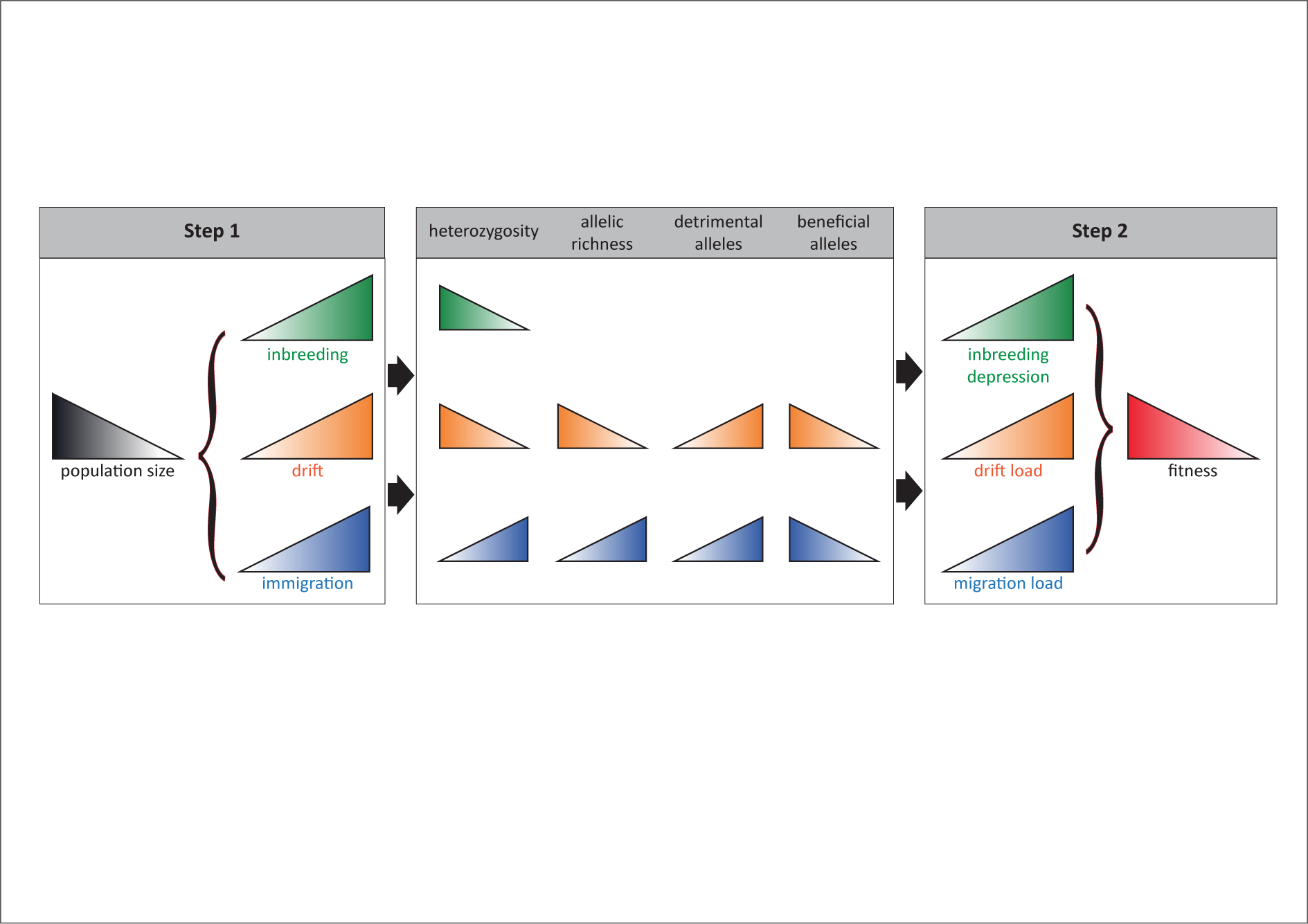
Scenarios underlying a genetic Allee effect. During Step 1, a decrease in population size *causes* an increase in inbreeding, a change in the drift/selection balance in favor of drift, and/or a higher proportion of immigrants in the population (assuming that both selection and migration are density-independent). Various population genetic structure variables are affected: heterozygosity, allelic richness, and the frequencies of detrimental and beneficial alleles. During step 2, these changes in genetic variation *cause* a decrease in the value of fitness components through inbreeding depression, drift load or migration load.

### Drift load

Here, we define drift as a change in allele frequencies due to the random sampling of alleles from one generation to the next (Wright 1969), and drift load as the decrease in the mean fitness of a population due to drift (Charlesworth et al. 1993, Whitlock 2000). Genetic drift is a consequence of finite population size and, like other random processes, such as demographic stochasticity, the manifestations of genetic drift are more evident in small populations (Fig. 2; Gabriel et al. 1991, Gabriel and Bürger 1994, Lynch et al. 1995a, Oostermeijer et al. 2003, Frankham 2005, Grueber et al. 2013). The balance between drift and selection determines allelic and genotypic frequencies. Selection is a deterministic process whereby the frequency of an allele (or a genotype) changes across generations according to its effect on fitness: beneficial alleles increase in frequency, whereas deleterious alleles decrease in frequency. Unlike drift, the intensity of which increases with decreasing population size, selection does not vary in intensity as a function of population size (Fig. 2), *i.e.*, we do not consider density-dependent selection here. Hence, in small populations, drift intensity often overwhelms selection, so that any changes in the number and frequencies of alleles result principally from drift rather than selection (Kimura 1983, Whitlock 2000).

The change in the selection/drift balance at small population size modifies the frequency of beneficial alleles, allelic richness and the heterozygosity of populations, driving the first step of the genetic Allee effect (Fig. 2). The exact nature of the change in genetic structure depends on the type of selection at work. When drift overwhelms negative selection, mildly deleterious alleles increase in frequency (Lanfear et al. 2014). Mildly deleterious alleles sometimes become fixed and constitute the drift load (Whitlock 2000, Glémin 2003). The corollary is that some beneficial alleles, despite being under positive selection, decrease in frequency or even disappear (Lanfear et al. 2014). The value of fitness components decreases if these components are influenced by loci that fix detrimental alleles and lose beneficial ones (Step 2). Variations in larval body size in the European common frog *Rana temporaria* provide an interesting example, with drift stronger than selection in small populations and weaker than selection in large populations (Step 1). Consequently, small populations have a higher drift load on larval body size, and display lower values for this fitness component (Table 1B; Johansson et al. 2007). Balancing selection also results in drift load. Balancing selection maintains several alleles at a locus under selection. Balancing selection can be due to overdominance and negative frequency-dependent selection. In small populations, in which drift overwhelms balancing selection, rare alleles can be lost (Zayed and Packer 2005, Levin et al. 2009, Eimes et al. 2011) and allele frequencies move away from the optimum value (Step 1), decreasing the values of the fitness components they influence (Step 2). For instance, plant self-incompatibility loci undergo balancing selection, triggering a genetic Allee effect. This has been shown in the rare plant *Brassica insularis,* in which smaller populations have fewer alleles at the self-incompatibility locus and, thus, lower rates of fruit set (Glémin et al. 2008).

### Migration load

Migration load is defined as the decrease in the mean fitness of a population due to the immigration of maladapted alleles and outbreeding depression (e.g. Bolnick and Nosil 2007). With effective dispersal, immigrants from a source population can bring new alleles into a sink population. Assuming a constant number of dispersers, the proportion of new alleles in the gene pool is higher in small sink populations than in large ones (Fig. 2), so the change in genetic structure depends on population size, fulfilling the conditions for the first step of the genetic Allee effect (Fig. 1). A migration load can occur as a result of local maladaptation and/or outbreeding depression (Ronce and Kirkpatrick 2001, Lenormand 2002) due to underdominance or deleterious epistatic interactions (Edmands 2002). The overrepresentation of new alleles in small populations results in a higher migration load in small populations, satisfying the conditions for the second step of the genetic Allee effect (Fig. 2). We were able to identify a single published example of a genetic Allee effect due to migration load (Table 1C). Populations of *Eucalyptus aggregata* can hybridize with closely related species of eucalypts; the hybridization rate increases with decreasing population size, and germination rates and seedling survival are lower in populations with a higher proportion of hybrids (Field et al. 2008).

### Combined mechanisms

As already reported for ecological component Allee effects (Berec et al. 2007), several genetic Allee effects may occur simultaneously in the same population. A good example is provided by the concomitant influences inbreeding depression, drift load, and loss of self-incompatibility alleles on small populations of the buttercup *Ranunculus reptans* (Willi et al. 2005). Genetic Allee effects can also act in conjunction with ecological Allee effects, as shown in experimental populations of the self-incompatible plants *Raphanus sativus* and *Lolium multiflorum* (Elam et al. 2007, Firestone and Jasieniuk 2013).

## How can a genetic Allee effect be detected?

The detection of a genetic Allee effect requires investigations of each of the two successive steps: (1) a decrease in population size *causing* a change in the within-population genetic structure, and (2) a decrease in a fitness component *caused* by this change. For demonstration of the causal relationship underlying step 2, two confounding effects must be avoided: environmental effects and ecological Allee effects. It is possible to control for environmental effects by using experimental designs involving constant environments (e.g., Charlesworth 2006, Oakley and Winn 2012) or extracting fitness responses to environmental variability before analyzing the effect of population genetic structure (e.g., Saccheri et al. 1998, Hanski and Saccheri 2006). However, controlling for environmental variability may nevertheless fail to separate genetic and ecological Allee effects, and some previously described ecological Allee effects may actually turn out to be genetic Allee effects. For instance, mating failures in small populations may result from a shortage of genetically compatible mates rather than a decrease in the frequency of encounters with conspecifics (Amos et al. 2001, Moller and Legendre 2001, Kokko and Rankin 2006). Only by experimental manipulations of population size and genetic variation with factorial designs can the genetic and ecological underpinnings of Allee effects be correctly disentangled. For example, genetic variation can be manipulated by modifying the number of founding families, whilst keeping census size constant (e.g., Elam et al. 2007, Hufbauer et al. 2013, Szücs et al. 2014).

Three main mechanisms underlie the second step of genetic Allee effects: inbreeding depression, drift load, and migration load. Inbreeding depression occurs when individuals with higher inbreeding coefficients have lower fitness. It can be assessed by studying individuals from populations with known inbreeding coefficients or offspring from controlled inbred and outbred crosses (e.g., Elam et al. 2007, Hufbauer et al. 2013). Drift load is detected when between-population crosses result in heterosis, and if this effect is stronger in populations with lower heterozygosity or allelic richness (Escobar et al. 2008, Oakley and Winn 2012). The respective strengths of drift and selection can be estimated by comparing *F_st_* and *Q_st_* between pairs of populations, under certain conditions: (i) comparison of several groups of populations, with similar population sizes within groups and different population sizes between groups (*F_st_* and *Q_st_* are calculated between all pairs of populations within groups); (ii) the component of fitness measured is a quantitative trait, with a continuous distribution, influenced by multiple loci with small effects. If estimates of *F_st_* and *Q_st_* are similar for a quantitative fitness component, then drift is probably stronger than selection for this component. A positive relationship between the value of this fitness component and the difference between *Q_st_* and *F_st_* may indicate a relationship between genetic variation and fitness (Oakley and Winn 2012). In the particular case of drift load involving plant self-incompatibility and balancing selection, the role of self-incompatibility in the relationship between genetic structure and fitness can be investigated by determining the proportion of incompatible crosses for which plants received pollen but produced no seeds (Fischer et al. 2003, Willi et al. 2005, Glémin et al. 2008). Finally, the demonstration that immigrants and/or hybrids have lower fitness components than residents is indicative of the presence of a migration load.

## At the core of the extinction vortex

A central concept in the biology of small populations is the extinction vortex, defined as a positive feedback between environmental, demographic and genetic factors, reinforcing one another in a downward spiral until the last individual has disappeared (the F-vortex; Gilpin and Soulé 1986). Meta-analyses of small populations suggest that extinction vortices do occur in declining populations (Fagan and Holmes 2006), and a body of models and data support the significant roles of genetic factors (Frankham and Ralls 1998, Tanaka 2000, Spielman et al. 2004, Frankham 2005). Nonetheless, despite its importance for the management of threatened species, the extinction vortex suffers from the rarity of clear demonstrations.

Formalizing the causal relations between population size, genetic structure and fitness, via the genetic Allee effect, should strengthen the approach of extinction vortices (Fig. 3). The rationale is that thinking in terms of Allee effects makes it possible to capitalize on the known duality of component and demographic Allee effects, thereby bridging the gap between individual fitness and population growth (Fauvergue 2013). Here, we have defined the genetic Allee effect as a component Allee effect, that is, an effect of population size on fitness, triggered by genetic factors. Like any other component Allee effect, the genetic Allee effect can lower mean individual fitness in small populations, thereby causing a decrease in the population growth rate, i.e., a demographic Allee effect (Lennartsson 2002; Fig. 3, Angulo et al. 2007). In the case of strong demographic Allee effect with negative growth rate, or if combined with detrimental environmental factors, the genetic Allee effect can drive small populations to extinction and is thus at the core of an extinction vortex.

**Figure 3.**
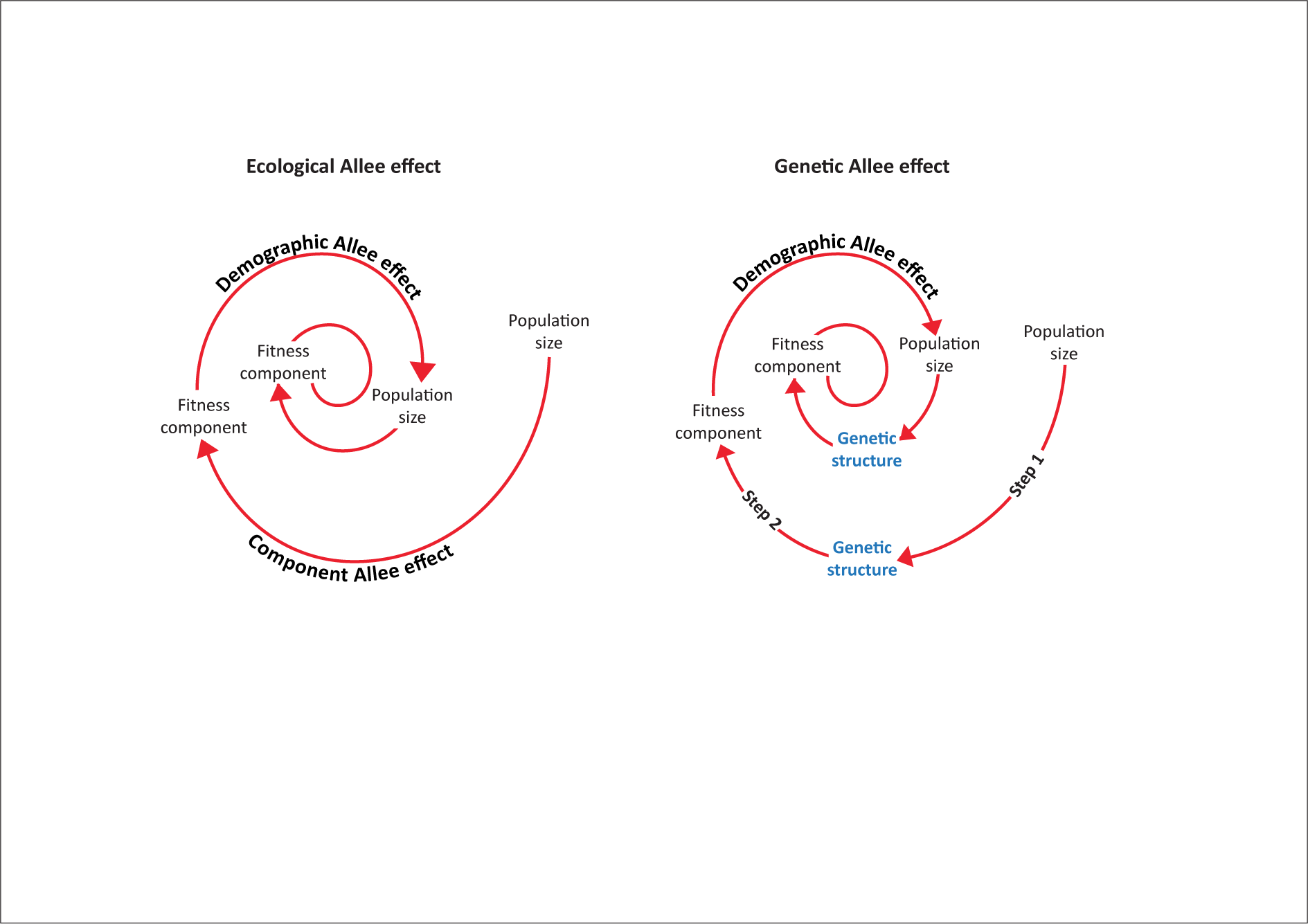
Extinction vortices driven by Allee effects. In ecological Allee effects, a decrease in population size directly causes a decrease in a component of individual fitness, which may in turn yield a decrease in population growth rate. A genetic Allee effect is a two-step process. A decrease in population size induces a change in the genetic variation of the population via inbreeding, drift/selection balance or migration (Step 1). This change yields inbreeding depression, drift load or migration load, causing a decrease in the value of fitness components (Step 2). Both ecological and genetic component Allee effects can produce a demographic Allee effect, which, if strong, can generate an extinction vortex.

A canonical example is provided by research on the Glanville fritillary butterfly, *Melitaea cinxia*. In this species, small subpopulations are less heterozygous than large subpopulations, and less heterozygous females produce larvae with lower survival rates (Saccheri et al. 1998). The Glanville fritillary is therefore subject to a genetic Allee effect. Controlled crosses in the laboratory and in the field have confirmed the existence of inbreeding depression in this species (Nieminen et al. 2001). The combination of inbreeding depression with poor resource availability causes the extinction of small subpopulations of *M. cinxia*, as shown by the higher probability of extinction for smaller and less heterozygous populations living in less favorable environmental conditions (Saccheri et al. 1998). A genetic Allee effect also impacts demography in the plant *Clarkia pulchella*. Using an experimental approach, Newman and Pilson (1997) showed that populations founded by genetically related seedlings were subject to inbreeding depression and/or drift load and therefore had a lower growth rate than populations founded by unrelated seedlings. In several other species, genetically eroded populations have been shown to have a lower growth rate (Fauvergue and Hopper 2009, Markert et al. 2010, Wennersten et al. 2012, Turcotte et al. 2013, Vercken et al. 2013). Fitness components were not measured in these studies, but the dynamics observed may reflect genetic Allee effects.

## Genetic versus ecological Allee effects

### Temporal issues

Genetic and ecological Allee effects are both component Allee effects capable of generating demographic Allee effects, but they differ in several ways, including the time scale. Genetic Allee effects are characterized by a time lag between the change in population size and its consequences for mean individual fitness, whereas an ecological Allee effect may occur as soon as population size decreases (*e.g.*, mating success decreases together with population density). The two steps of a genetic Allee effect may not occur over similar time scales: a decrease in population size may modify the genetic structure of the population over tens or thousands of generations, but fitness consequences may appear in the generation following the change in genetic structure (Amos and Balmford 2001, but see Hufbauer et al. 2013). Conversely, if population size increases, ecological Allee effects should die down almost immediately, whereas genetic Allee effects will not. For instance, after several generations of genetic drift, a population cannot recover its initial level of genetic variation unless new alleles arise by mutation or migration. This implies that a population can be rescued from a demographic threat, but, if the underlying genetic variation is not also restored, it may still suffer from a genetic Allee effect and be prone to extinction (what has been referred to as an extinction debt; Vercken et al. 2013).

### Two steps for the genetic Allee effect

The two steps of genetic Allee effects are idiosyncratic: each step can occur independently of the other, but the occurrence of the two steps in succession is necessary to create (and demonstrate) a genetic Allee effect. The first step, a change in genetic structure caused by a decrease in population size, can occur with no decrease in mean fitness (Step 1, but not Step 2). For instance, inbred individuals do not necessarily suffer from inbreeding depression. Indeed, although counterintuitive at first glance, inbreeding can even result in the purging of deleterious alleles, leading to an increase in mean fitness (Glémin 2003, Facon et al. 2011). Similarly, drift can randomly eliminate detrimental alleles and fix beneficial ones, and immigration in small populations can lower the negative impact of drift and inbreeding by bringing new adapted alleles and increasing heterozygosity. The second step, in which a change in the genetic structure of a population triggers a decrease in a component of fitness, is not necessarily induced by the first step, because genetic structure can change independently of population size, under the influences of mutation, migration or the mating system.

### Evolutionary consequences

The effects on individual fitness of ecological Allee effects could act as a selective force, driving the evolution of adaptations mitigating sensitivity to population size (Courchamp et al. 2008). For instance, long-range volatile sex pheromones may prevent mating failures (Fauvergue et al. 2007). The same reasoning applies to genetic Allee effects too. Inbreeding depression may have shaped the evolution of dispersal, inbreeding avoidance and self-incompatibility systems (Penn and Potts 1999, Perrin and Mazalov 2000).

## Conclusion: why is all this important?

The focus on genetic Allee effects does not simply add a new semantic layer to processes that have been thoroughly discussed in the last few decades. On the contrary, it is essential, for several reasons.

First, although a genetic Allee effect has been mentioned in a few articles (starting with Fischer et al. 2000), no formal definition has ever been provided. We show here that a genetic Allee effect is not merely an Allee effect underpinned by genetic mechanisms. The formal definition we provide here includes two successive steps: a decrease in population size modifying the within-population genetic structure, in turn causing a decrease in the mean value of a component of individual fitness. The requirement of these two steps differentiates genetic Allee effects from ecological Allee effects. Genetic Allee effects are unique in terms of their potentially long time scale (particularly for the change in genetic structure in response to a decrease in population size) and their entropy (by contrast to ecological Allee effects, a population cannot recover instantly from a genetic Allee effect). According to the formal definition proposed here, some reported findings refer to a genetic Allee effect (including from our own research; Fauvergue and Hopper 2009, Vercken et al. 2013), but without providing explicit evidence for such an effect. Alternatively, some specific types of Allee effect, such as the so-called S-Allee effect (Wagenius et al. 2007, Hoebee et al. 2012, Busch et al. 2014), may be seen as genetic Allee effects. The provision of a clear definition of the genetic Allee effect should prompt a better examination of the genetic mechanisms affecting fitness and demography in small populations. We identified only 15 demonstrations of this particular type of component Allee effect in the literature, but we predict that widespread evidence of its existence will be obtained if the methods we propose are used to identify genetic Allee effects and to distinguish them from ecological Allee effects occurring in the same populations.

Second, a definition of the genetic Allee effect should promote a more comprehensive approach to the biology of small populations. The integration of demography and genetics has long been recognized as an important endeavor, but tangible progress is still required, particularly for the management of declining and/or bottlenecked populations (e.g., Robert et al. 2007, Fauvergue et al. 2012). Indeed, early developments in population viability analyses clearly assumed a feedback between demographic and genetic processes at the core of extinction vortices (Gilpin and Soulé 1986, Caughley 1994) and, more recently, the concept of evolutionary rescue has emerged as a race between demographic and evolutionary processes (Bell and Gonzalez 2009, Gonzalez et al. 2013). However, these concepts have generally been restricted to the analysis of demographic and environmental stochasticity, resulting in the neglect of the Allee effect as an important mechanism for small populations. Conversely, the Allee effect is attracting the attention of an increasing number of scientists with a general interest in the biology of small populations and extinction vortices. However, despite widespread evidence for the significant role of genetic processes, it is only very recently that theoretical works have started to combine Allee effects with genetic processes (Kanarek and Webb 2010, Roques et al. 2012, Shaw and Kokko 2014, Wittmann et al. 2014a, b, Kanarek et al. 2015). The genetic Allee effect, as defined here, is at the very intersection of demography and genetics and should serve as a unifying paradigm. In the long term, the genetic Allee effect should contribute to improving the dialog between genetics and demography.

## Acknowledgments

This work was supported by the *Agence Nationale de la Recherche* (Sextinction project ANR-2010-BLAN-1717 and RARE project ANR-2009-PEXT-01001) and the *Fédération de Recherche sur la Biodiversité* (VORTEX project APP-IN-2009-052). We thank four anonymous reviewers for valuable insights on previous versions of this manuscript, and Nathan Fauvergue for fresh thoughts.

